# Alpha oscillations during incidental encoding predict subsequent memory for new “foil” information

**DOI:** 10.1101/141648

**Authors:** David A. Vogelsang, Matthias Gruber, Zara M. Bergström, Charan Ranganath, Jon S. Simons

## Abstract

People can employ adaptive strategies to increase the likelihood that previously encoded information will be successfully retrieved. One such strategy is to constrain retrieval towards relevant information by re-implementing the neurocognitive processes that were engaged during encoding. Using electroencephalography (EEG), we examined the temporal dynamics with which constraining retrieval towards semantic versus non-semantic information affects the processing of new “foil” information encountered during a memory test. Time-frequency analysis of EEG data acquired during an initial study phase revealed that semantic compared to non-semantic processing was associated with alpha decreases in a left frontal electrode cluster from around 600ms after stimulus onset. Successful encoding of semantic versus non-semantic foils during a subsequent memory test was related to decreases in alpha oscillatory activity in the same left frontal electrode cluster, which emerged relatively late in the trial at around 1000–1600ms after stimulus onset. Across subjects, left frontal alpha power elicited by semantic processing during the study phase correlated significantly with left frontal alpha power associated with semantic foil encoding during the memory test. Furthermore, larger left frontal alpha power decreases elicited by semantic foil encoding during the memory test predicted better subsequent semantic foil recognition in an additional surprise foil memory test. These findings indicate that constraining retrieval towards semantic information involves re-implementing semantic encoding operations that are mediated by alpha oscillations, and that such re-implementation occurs at a late stage of memory retrieval perhaps reflecting additional monitoring processes.

## Introduction

Memory retrieval often requires goal-directed control processes in order to optimize retrieval success. One possible strategy people use to facilitate memory retrieval is to re-implement the neurocognitive processes that were involved during encoding (Rugg et al., 2008). This idea has been manifested in the Transfer Appropriate Processing Framework, which states that the likelihood of retrieval success is dependent on the overlap between encoding and retrieval operations (Morris et al. 1977; Roediger et al., 1989). Accordingly, the most effective encoding strategy depends on the specific conditions at retrieval and, conversely, what constitutes an optimal retrieval strategy depends on the conditions under which the information was encoded (Rugg et al., 2008). In line with this principle, previous behavioral studies have shown that recognition memory is enhanced when mental operations at encoding are recapitulated during a subsequent memory test (Dewhurst & Brandt, 2007; Morris et al., 1977).

Jacoby and colleagues (2005a) developed a behavioral “memory for foils” paradigm that enabled such encoding-retrieval overlap to be investigated more directly. In an initial study phase (phase 1 of the paradigm), participants studied nouns in two separate blocks, one of which involved a semantic task (pleasant/unpleasant judgment) and the other a non-semantic task (letter judgment). In a subsequent recognition memory test (phase 2), studied and non-studied words were intermixed and participants undertook blocks in which they judged whether they had previously encountered the words in the pleasantness judgment task or whether they were new (the memory test for semantically encoded words), or judged whether they had previously seen the words in the letter judgment task or whether they were new (the memory test for non-semantically encoded words). Of special interest were the new words (so called ‘foils’) in the semantic and non-semantic memory test blocks. The semantic and non-semantic foils were subsequently mixed together with completely new words in a final foil recognition test (phase 3) in which participants were again instructed to make an old/new judgment, this time about whether the words had been encountered at any time during the experiment or were completely novel. Jacoby et al. found that the “foil” words were differentially memorable depending on the type of test in which they had been originally encountered: recognition memory was significantly higher for semantic compared to non-semantic foils. Because semantic encoding typically leads to more accurate memory compared to non-semantic encoding, this “foil effect” implies that participants strategically orient their retrieval towards a semantic processing mode when attempting to retrieve semantic encoded information, and a non-semantic processing mode when retrieving non-semantic information, resulting in better incidental encoding of semantic compared to non-semantic foils. Jacoby and colleagues interpreted this foil finding in light of the transfer appropriate processing principle by emphasizing the importance of the overlap in study-test operations for optimizing retrieval success (see also Alban and Kelley, 2012; Danckert et al., 2011; Gray & Gallo, 2015; Halamish et al., 2012; Kantner and Lindsay, 2013; Marsh et al., 2009; Zawazka et al., 2017).

Recently, we collected functional magnetic resonance imaging (fMRI) data in a “memory for foils” paradigm and applied subsequent memory analysis (also known as “difference due to memory” or “DM effect”) to study the neural mechanisms underlying the enhanced encoding of foils in a semantic compared to non-semantic recognition test. The results revealed that the left inferior frontal gyrus (LIFG) exhibited significantly greater subsequent memory effects for semantic compared to non-semantic foils (Vogelsang et al., 2016). A conjunction analysis revealed significant overlap in activity between semantic processing in the initial study phase and semantic foil encoding during the first memory retrieval test in the LIFG; however, this overlap in activation was not observed for the non-semantic condition. The LIFG has previously been associated with semantic processing and semantic encoding strategies across many studies (Fletcher et al., 2003; Kim, 2011; Poldrack et al., 1999; Wagner et al., 1998). Together with the behavioral result that semantic foils were recognized more accurately than non-semantic foils on the final surprise foil recognition test, these neuroimaging data support the hypothesis that directing retrieval towards new semantic versus non-semantic information leads to the recruitment of distinct neural mechanisms that are predictive of subsequent memory (Vogelsang et al., 2016).

A key element of the foil paradigm is the proposal that retrieval is strategically oriented towards the relevant processing mode to facilitate memory search before information is retrieved. This account suggests that the neural mechanisms that underlie retrieval orientation are engaged shortly after a memory cue is encountered in order to guide retrieval attempts (a form of “front-end control”, Jacoby et al., 2005b). Alternatively, strategic control processes can also be recruited later on in the trial when retrieval attempts have failed or have produced ambiguous information and additional monitoring or verification is required (a form of “backend control”, Halamish et al., 2012; or “late correction strategy”, Jacoby et al. 1999). Previous fMRI research was unable to distinguish these accounts (Vogelsang et al., 2016) because the low temporal resolution of the blood-oxygen-level dependent (BOLD) precludes investigation of at which stage of retrieval (early versus late) LIFG activity is reinstated for semantic compared to non-semantic foils. Therefore, in the current study, we recorded electroencephalography (EEG) oscillations during performance of the “memory for foils” paradigm. The fine-grained temporal resolution of neural oscillations can provide more information with regard to the question of *when* the neural activity associated with initial encoding operations during a study phase re-occur during the incidental encoding of foils in a subsequent recognition test.

Neural oscillations and their relationship with memory functions have gained considerable interest in recent years (Fell & Axmacher, 2011). In the memory encoding literature, there is evidence that a decrease in alpha power might be related to semantic processing (Bastiaansen et al., 2005; Hanslmayr et al., 2009; Hanslmayr & Staudigl, 2014; Zion-Golumbic et al., 2009; for review see Klimesch, 1999). For example, Hanslmayr and colleagues (2009) contrasted deep semantic encoding with shallow non-semantic encoding, and found power decreases in alpha (and beta) frequency bands that were related to successful semantic encoding. Fellner and colleagues (2013) showed that alpha and beta decreases predicted subsequent memory in a semantic condition, but not in a non-semantic but still highly efficient encoding condition (in this case a survival processing task), thereby suggesting that alpha decreases are likely a reflection of semantic processing in particular, rather than of efficient encoding strategies in general. Furthermore, alpha decreases have been observed over left frontal electrodes in tasks requiring high semantic processing demands (Hanslmayr & Staudigl, 2014; Klimesch, 1999; Wang et al., 2012;), but have also been associated with subsequent memory effects (Klimesch et al., 1997), consistent with the idea that the left prefrontal cortex is important for successful encoding (Vogelsang et al., 2016; Wagner et al., 1998).

The main aim of the present experiment was to investigate the temporal dynamics of EEG oscillations associated with encoding of new “foil” words during a memory retrieval test. We focused our analysis on alpha EEG frequencies (8–10Hz) because previous research has shown that alpha plays a role in both semantic processing (Bakker et al., 2015) and subsequent memory effects (Hanslmayr et al., 2009). We first examined alpha power associated with semantic versus non-semantic processing during the initial study phase. We then investigated whether alpha power differences were again observed during successful encoding of semantic versus non-semantic foils in the first memory test, which would support the hypothesis that the incidental encoding of foils in a memory test involves the re-implementation of the neurocognitive processes that were involved during initial encoding (Bergström et al., 2015; Jacoby et al., 2005a; Jacoby et al. 2005b; Vogelsang et al., 2016). Most importantly, the high temporal resolution of EEG oscillations allowed us to examine whether alpha reinstatement during foil encoding occurred early or late in the trial, which we hypothesized would indicate that participants used “front end” or “back end” control strategies, respectively. We also tested whether those individuals who showed the largest alpha power differences during semantic versus non-semantic processing in the study phase also showed the largest alpha power differences during semantic encoding of foils in the retrieval test, which would support the hypothesis that the alpha effects during study and test were functionally related. Finally, we tested the hypothesis that if alpha frequencies mediate semantic foil encoding during the first recognition test, then individuals who showed larger alpha differences for successfully encoded foils during the first test should also exhibit better foil recognition performance in the final foil recognition test.

## Methods

### Participants

Fifty-three right handed healthy English native speakers with normal or corrected to normal vision participated in this experiment. Written informed consent was obtained before commencement of the experiment and all participants received £15 for their participation. Data from 17 participants were excluded because they did not produce enough trials of each type for the subsequent memory analysis (see “Time-Frequency Analysis” below for details). Additionally, data from two participants were excluded because of excessively noisy EEG data. The final dataset thus consisted of 34 participants (21 female, mean age = 21.9 years, range 18–33). The study was approved by the University of Cambridge Psychology Research Ethics Committee.

### Materials

The stimuli consisted of 552 nouns (e.g. “book”) derived from the MRC psycholinguistic database (Wilson, 1988; also used in Vogelsang et al., 2016). The 552 words were split into 6 lists that were matched for concreteness, familiarity, Kucera-Francis Frequency, word length and number of syllables, and we counterbalanced the assignment of lists to the experimental conditions across participants.

### Procedure

Participants were fitted with an EEG cap and were seated in a sound and light attenuated room. Participants completed three phases: 1) A study phase (henceforth referred to as “phase 1”), 2) An initial memory test (henceforth referred to as “phase 2”), and 3) A final surprise memory test that assessed foil recognition (henceforth referred to as “phase 3”). Throughout all phases, participants provided their responses on a button box with either their left or right hand (counterbalanced across participants).

Phase 1 consisted of two separate incidental encoding blocks during which participants were instructed to make a simple judgment. In a semantic judgment study block, participants made a pleasantness judgment (“Is this word pleasant?”). In a non-semantic study block, participants made a letter judgment (“Is there a letter O or U in the word?”). Each trial in the study phase started with a 500ms fixation cross followed by the stimulus that was presented in the center of the screen for 2000ms. Both the semantic and non-semantic judgment blocks consisted of 92 trials each. The order of semantic and non-semantic judgment blocks was counterbalanced across participants. Participants were instructed to respond while the words were on the screen.

In phase 2, participants’ memory for the stimuli encountered during phase 1 was assessed in an old/new recognition test, which aimed to manipulate whether participants oriented retrieval towards semantic or non-semantic information. In the semantic test phase, 92 old words from the semantic study phase were intermixed with 92 new words (semantic foils). In the non-semantic test phase, 92 old words from the non-semantic study phase were intermixed with 92 new words (non-semantic foils). For both blocks, participants were told in which specific phase 1 task any old words had been previously shown, in order to encourage them to engage different retrieval orientations for the two blocks. The order of test block (semantic and non-semantic) was counterbalanced across participants. Each test trial began with a 500ms fixation cross, followed by the presentation of the stimulus centrally on the screen for 2000ms. Participants were instructed to provide their response as to whether each word was old or new while the stimulus was still on the screen.

In the final phase 3, a surprise old/new foil recognition test (phase 3) was administered in which participants were asked to distinguish between the semantic and non-semantic foils and completely new words. Participants were instructed that they were “going to be presented with a word that is either old or new. ‘Old’ in this case means that you saw the word at some point earlier in the experiment in any study or test phase. ‘New’ words are words you have not seen at all in today’s experiment”. This foil recognition test consisted of 368 words (92 semantic foils, 92 non-semantic foils, and 184 completely new words, which were randomly intermixed). Each trial in the final foil recognition test began with a 500ms fixation cross followed by the stimulus presented centrally for 2000ms.

### EEG Recording and Preprocessing

EEG data was acquired during all phases of the experiment and was recorded with an Electrical Geodesic Netamps 200 system with a 128-channel HydroCel Geodesic Sensor Net (GSN 200, Tucker, 1993). The recorded EEG data was referenced to Cz and was filtered with a bandwidth of 0.01–70 Hz (250 Hz sampling rate). The EEG data was analyzed in EEGLab 13 (Delorme & Makeig, 2004). The continuous EEG data from the study phase and first retrieval test was re-referenced to an average mastoid reference, and high pass filtered with a cut-off of 0.5Hz (two-way least-squares finite impulse response filter) and the continuous data were divided into epochs ranging from -500ms before cue onset until 2000ms thereafter. Artifact correction was applied using extended info-max Independent Component Analysis (ICA; Bell & Sejnowski, 1995, in Delorme & Makeig, 2004) using Runica from the EEGLab toolbox, with default mode training parameters (Delorme & Makeig, 2004). Independent components reflecting eye movements and other sources of noise were identified by visual inspection of component scalp topographies, time courses, and activation spectra and were discarded from the data by back-projecting all but these components to the data space. Trials that still contained artifacts after running ICA correction, were removed after visual inspection, resulting in only 5–10% of the trials being excluded.

### Time-Frequency Analysis

Time-frequency analysis in EEGLab was applied using Morlet wavelets (Percival & Walden, 1993) with 6 cycles in a frequency range of 4–30Hz, with steps of 1Hz between each wavelet center frequency. These wavelets were applied in a sliding window with 20ms increments in the 0–2000ms interval. In order to minimize edge effects (distortions that occur at the edge of the time window of analysis), we concatenated mirrored (i.e. time inverted) segments at the left and right edge of the original epochs. We then performed the wavelet transform on these extended epochs, and discarded the concatenated segments from the final analysis (for detailed explanation of this “reflection approach” see Cohen, 2014; and see Fell et al., 2011, for example of a paper using this approach). Baseline correction in the frequency domain was applied for each epoch by subtracting the mean voltage of 0–200ms before stimulus onset (see for similar procedures Hsieh et al., 2011).

In order to identify the neural oscillations associated with sematic and non-semantic processing, we first examined the power spectra of epoched data from phase 1. For each of the 34 participants, EEG data during the study phase were binned according to the type of processing (semantic vs. non-semantic). In this way, we could isolate the EEG frequencies that were elicited by semantic and non-semantic processing in order to later examine whether these frequencies were reinstated during the encoding of foils in the first test phase (phase 2). Mean trial numbers were the following: semantic study mean = 90, range 46–92; non-semantic study mean = 92, range 88–92.

To analyze the neural oscillations during phase 2, we binned the EEG data for each participant according to condition (semantic vs. non-semantic) and subsequent memory (remembered vs. forgotten). Mean trial numbers for each condition were: semantic foils remembered mean = 65, range 25–81; semantic foils forgotten mean = 25, range 12–67; non-semantic foils remembered mean = 55, range 15–79; and non-semantic foils forgotten mean =35, range 13–77.

Time-frequency analysis was conducted on EEG that was averaged within nine electrode clusters (frontal vs. central vs. posterior; left vs. middle vs. right; see Figure 1), based on a previous study by Hsieh et al. (2011). These clusters included the following: left frontal cluster (channels 33, 24 and 26; equivalent to F3, F7, AF7), mid frontal cluster (channels 19, 11, 4; equivalent to Fz, F1, F2), right frontal cluster (channels 124, 2, 122; equivalent to F4, F8, AF8), left central cluster (channels 35, 36, 41; equivalent to C5, C3, T7), mid central cluster (channels 31, 55, 80; equivalent to Cz, C1, C2), right central cluster (channels 109, 104, 110; equivalent to C4, C6, T8), left posterior cluster (channels 52, 53, 60; equivalent to P3, P1, PO3), mid posterior cluster (channels 61, 62, 78; equivalent to CP1, Pz, CP2), and right posterior cluster (channels 85, 86, 92; equivalent to P2, P4, PO4).

**Figure 1.**
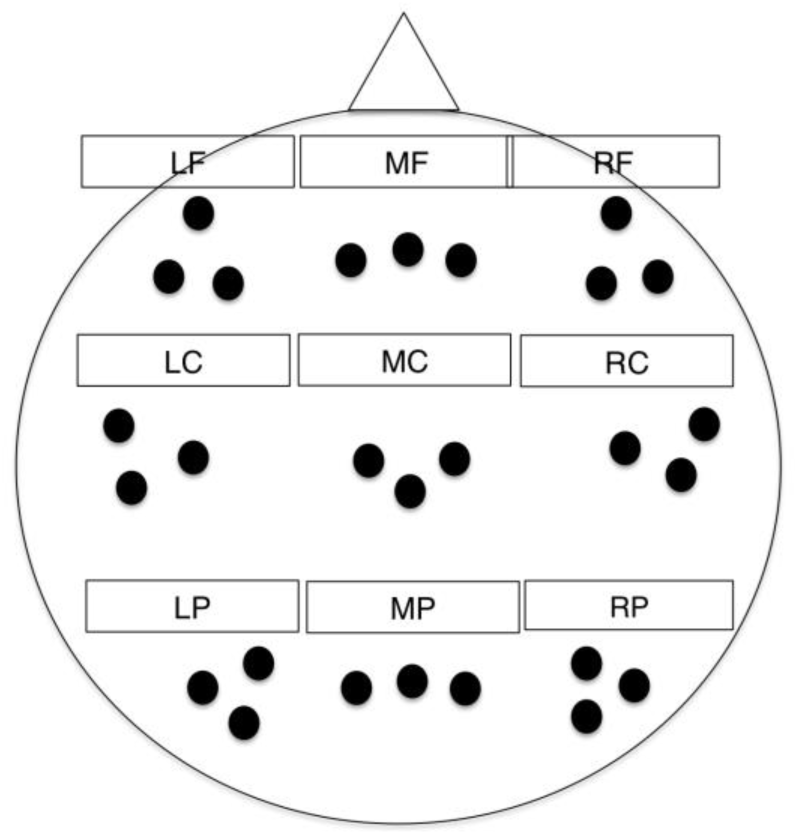
Locations of the electrode clusters, which included left frontal (LF), mid frontal (MF), right frontal (RF), left central (LC), mid central (MC), right central (RC), left posterior (LP), mid posterior (MP) and right posterior (RP).

### Statistical Analysis

Across trial permutation tests were conducted to test for significant effects in alpha power related to the semantic versus non-semantic processing in phase 1 and successful encoding of semantic versus non-semantic foils in phase 2. For both phases, the 2000ms epoch was split into time windows of 200ms each.

For phase 1, the permutation testing was conducted on the mean power alpha (8–10Hz) frequencies per condition for each time window and electrode cluster (see Gruber et al., 2013 for similar procedure). We first conducted two-tailed paired t-tests on the relevant electrode clusters comparing the two conditions. Secondly, the two conditions were then interchanged randomly for each subject and each randomization run, so that pseudo conditions were created in which systematic differences between the conditions were eliminated. This step was repeated 1000 times so that a null distribution of 1000 t-values were created. The two tails of the null t-distribution were used as critical t-values. Using an alpha level of 0.05 with 1000 permutations, we used the 25th and 975th values to represent the critical t-values and values below or higher than these values were considered to be significant effects. This permutation method was based on Blair and Karniski (1993; for similar approaches see Addante et al., 2011; Hanslmayr et al., 2009; Staudigl et al., 2010; Gruber et al., 2013). Significant effects in alpha power in consecutive time windows were collapsed and another permutation test was run on the extended time window. For simplicity, the results reported here are from the extended time windows (see for similar procedure Pastötter et al., 2011; Gruber et al., 2013).

For phase 2, we conducted the permutation testing only in those electrode clusters that showed a significant alpha effect in phase 1. Thus, the electrode clusters that showed a significant effect in phase 1 were taken as “electrode clusters of interest” for the analysis for phase 2 data, to examine alpha activity re-implementation during encoding of foils in the same electrode clusters that showed significant alpha effects in phase 1. To test our hypothesis that re-implementation of semantic processes facilitate successful encoding of foils, we focused on the interaction between condition (semantic vs. non-semantic) and subsequent memory in phase 2 (remembered vs. forgotten) by comparing the difference between remembered and forgotten semantic foils versus the difference between remembered and forgotten non-semantic foils. We also tested the simple effects of subsequent memory for semantic vs. non-semantic conditions separately. The rest of the permutation procedure was the same as for the phase 1 data.

### Data Availability

The data that support the findings of this study are available at the University of Cambridge data repository (http://doi.org/10.17863/CAM.9855).

## Results

### Behavioral Results

Recognition accuracy for phase 2 was calculated using the discrimination measure p(Hits)-p(False alarms) (Snodgrass & Corwin, 1988) and the results are presented in Table 1. Recognition memory for semantic trials was significantly more accurate compared to non-semantic trials (t(33)= 25.4, p < 0.001, 95% CI [0.47, 0.56], Cohen’s Dz = 4.4). Furthermore, RTs were faster for old semantic items compared to old non-semantic items (t(33) = 4.39, p < 0.001, 95% CI [49, 134], Cohen’s Dz = 0.75). Foils presented in the semantic condition were also endorsed as new significantly more quickly than foils presented in the non-semantic condition (t(33) = 2.23, p = .033, 95% CI [4, 84], Cohen’s Dz = 0.38).

**Table 1.** Accuracy (Hits and false alarms (FA)) and reaction time (for hits and correct rejections) for phase 2.

|  | **Hits** |  | **FA** |  | **RT(ms)** |  | *Correct Rejections (Mean)* | *Correct Rejections (SD)* |
| --- | --- | --- | --- | --- | --- | --- | --- | --- |
|  | *Mean* | *SD* | *Mean* | *SD* | *Hits (Mean)* | *Hits (SD)* |  |  |
| Semantic | 0.88 | 0.07 | 0.13 | 0.11 | 890 | 113 | 938 | 117 |
| Non-semantic | 0.50 | 0.15 | 0.26 | 0.13 | 981 | 138 | 982 | 161 |

The behavioral results of phase 3 are presented in Table 2. Note that we conducted the phase 3 analysis on accuracy scores (proportion correct) rather than Hits-FAs because in the final foil recognition test completely new items were intermixed with semantic and non-semantic foils, so a proper Hits-FAs analysis cannot be conducted here. In line with our main prediction, semantic foils were recognized significantly more accurately than non-semantic foils (t(35) = 5.18, p < 0.001, 95% CI [0.066, 0.15], Cohen’s Dz = 0.89), and significantly faster (t(33) = 3.42, p = .002, 95% CI [9, 37], Cohen’s Dz = 0.59). There was no significant difference in reaction time between non-semantic foils and new items (t(33) = 1.5, p = 0.15), however, RT was faster for recognizing semantic foils compared to new items (t(33) = 4.03, p < 0.001, 95% CI [17, 52], Cohen’s Dz = 0.69). These results replicate earlier findings of the “foil effect” obtained in previous studies (Jacoby et al., 2005a; Jacoby et al., 2005b; Bergström et al., 2015; Vogelsang et al., 2016).

**Table 2.** Accuracy (proportion correct) and reaction time for phase 3.

|  | **Accuracy** |  | **RT(ms)** |  |
| --- | --- | --- | --- | --- |
|  | Mean | SD | Mean | SD |
| Semantic Foils | 0.72 | 0.16 | 887 | 122 |
| Non-semantic Foils | 0.61 | 0.16 | 910 | 118 |
| New Items | 0.77 | 0.12 | 921 | 127 |

### Time-Frequency Results

#### Phase 1: Semantic versus Non-Semantic Processing

The time-frequency analysis of phase 1 focused on a direct comparison between all semantic and all non-semantic trials. The results are presented in Figure 2. Significant decreases in alpha power were observed over the left frontal electrode cluster between 600–1000ms after word onset (t(33) = −2.35, p = 0.025, 95% CI [−1.4, −0.1], Cohen’s Dz = 0.44). Furthermore, significant power decreases in alpha were also observed between 600–1600ms after word onset over mid and right posterior electrode clusters: mid posterior (t(33) = −2.57, p = 0.015, 95% CI [−1.11, −0.13], Cohen’s Dz = 0.44) and right posterior (t(33) = −2.19, p = 0.035, 95% CI [−0.90, −0.03], Cohen’s Dz = 0.38). The time course of the alpha power changes in the left frontal electrode cluster is presented in Figure 3.

**Figure 2.**
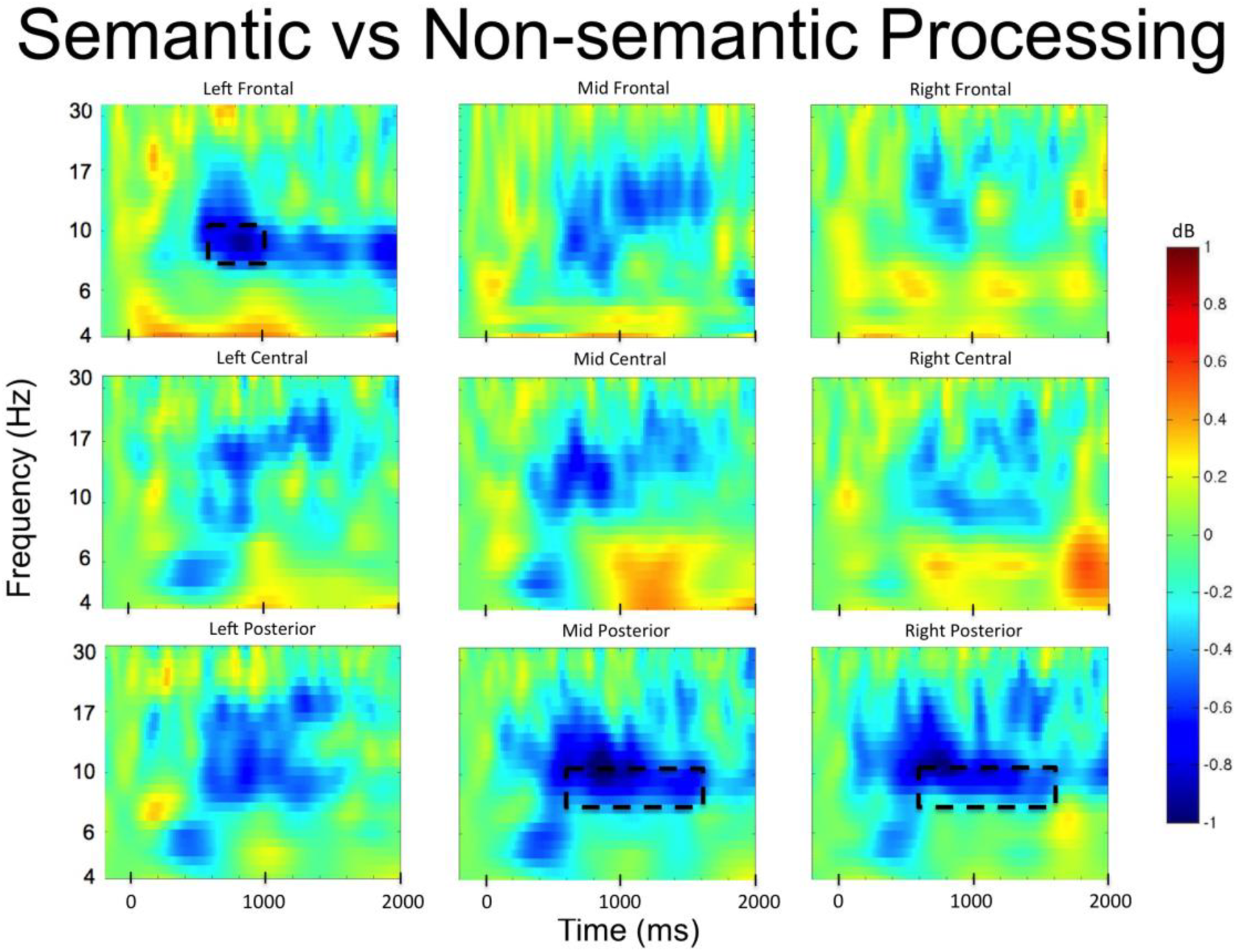
Time-frequency plots for semantic versus non-semantic processing in the study phase. Significant decreases in alpha frequencies were observed in left frontal and mid and right posterior electrode sites.

**Figure 3.**
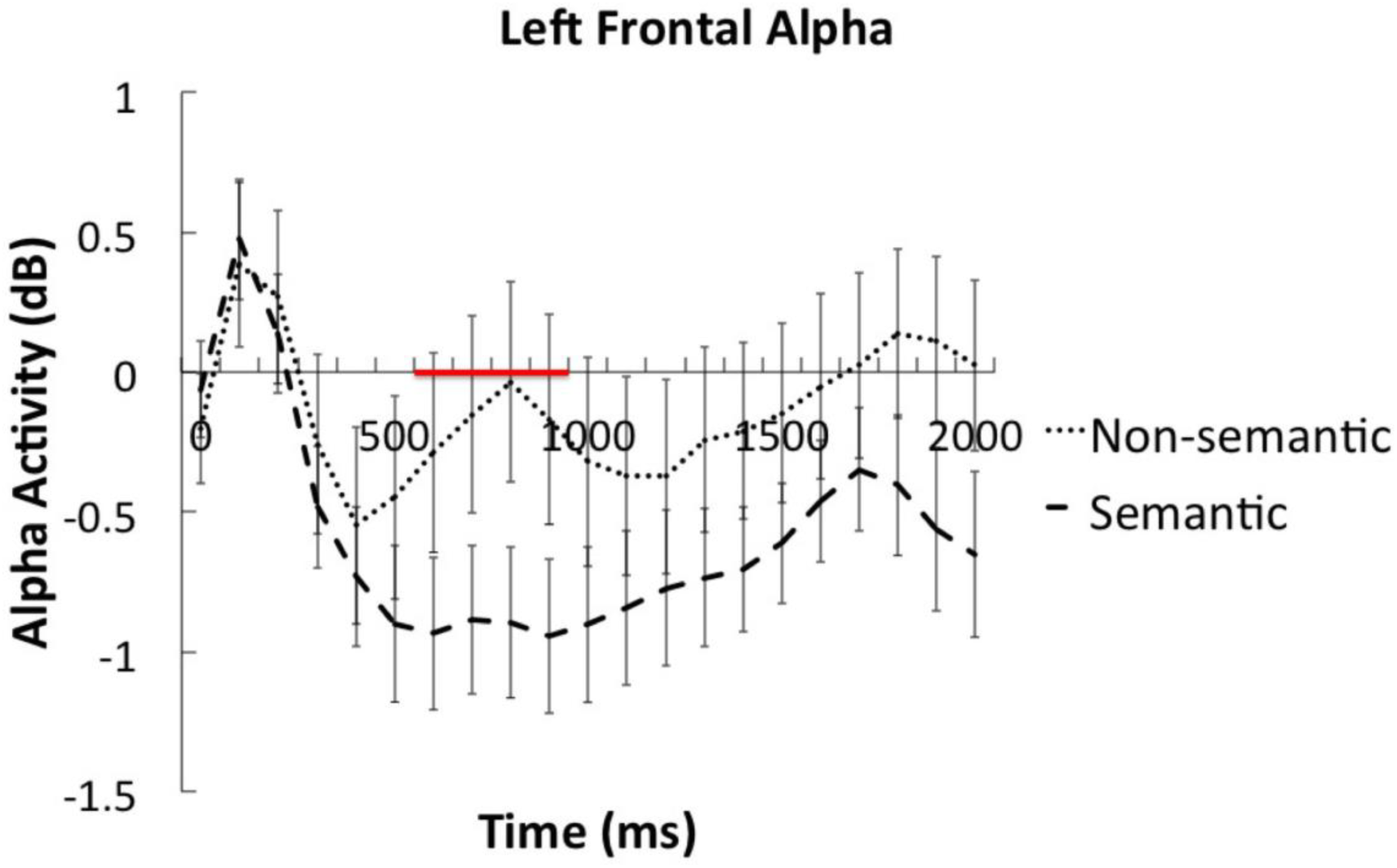
Average alpha (8–10Hz) frequency time courses (in decibel) for semantic and non-semantic processing in the left frontal cluster. Red line on x-axis represents significant time window.

### Subsequent Memory Effect for Foils in Phase 2

The second part of the time-frequency analysis focused on the temporal dynamics of subsequent memory effects (DM effect) for foils during phase 2 to investigate when alpha activity was reinstated in a way that facilitated encoding of semantic foils. The time-frequency plot of the interaction term (DM effect for semantic foils – DM effect for non-semantic foils) for all electrode clusters is presented in Figure 4. Since significant alpha effects in phase 1 were only observed in left frontal, mid and right posterior electrode clusters, only these three clusters were used to conduct the permutation analysis in phase 2, which allowed us to directly test the re-implementation hypothesis.

**Figure 4.**
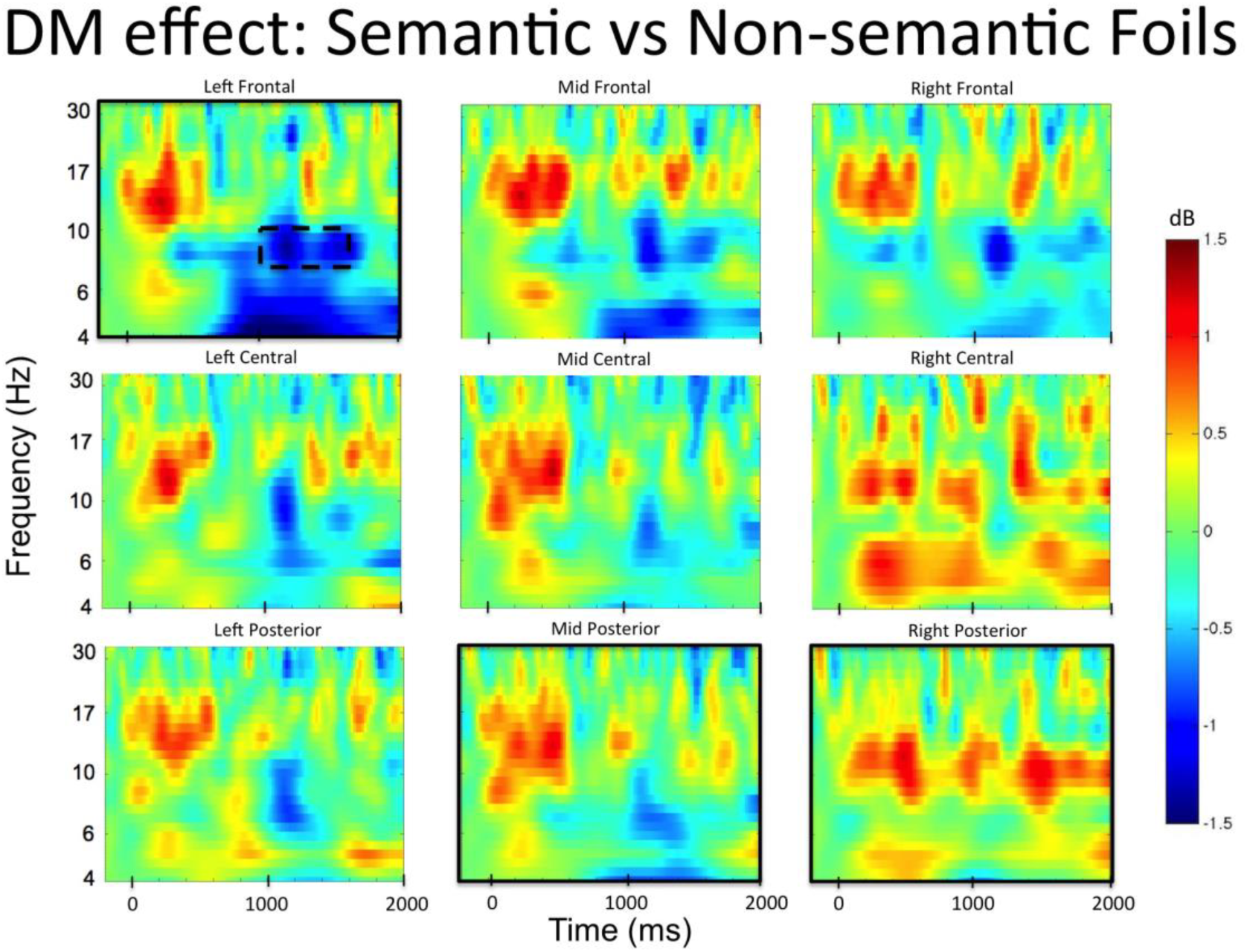
Time-frequency plots from the left frontal cluster illustrating EEG oscillation differences associated with successful encoding (DM effect) of semantic vs. non-semantic foils during the first memory test (phase 2). The plot represents the difference of semantic foils later remembered – forgotten versus non-semantic foils later remembered – forgotten. Permutation testing conducted on the left frontal, mid and right posterior electrode clusters (indicated by black squares) revealed a significan alpha decrease for subsequently remembered versus forgotten semantic versus non-semantic foils in the left frontal cluster (dashed box).

Out of the three electrode clusters used for the phase 2 analysis, only the left frontal electrode cluster showed a significant interaction in the alpha band between 1000–1600ms after word onset (t(33) = −2.31, p = 0.027, 95% CI [−1.80, −0.12], Cohen’s Dz = 0.40, see dashed box in Fig. 4). Time frequency plots comparing EEG oscillations associated with successful encoding of each type of foils separately are presented in Figure 5, and the time courses for alpha frequencies in the left frontal cluster for the semantic and non-semantic subsequent memory effect are presented in Figure 6. These comparisons indicated that the significant interaction arose because power differences between remembered and forgotten items were observed in the semantic but not in the non-semantic condition. For successful encoding of semantic foils, alpha in the 1000–1600ms time window (t(33) = −1.84, p = 0.074, 95% CI [-1.26, 0.06], Cohen’s Dz = 0.32) power approached significance depending on whether a word was later remembered or forgotten. However, no significant differences between remembered and forgotten words were observed for non-semantic foils (1000–1600ms alpha: t(33) = 1.64, p = 0.11, 95% CI [−0.086, 0.81], Cohen’s Dz = 0.28; and in fact power differences were numerically in the opposite directions in this condition). Thus, the subsequent memory effects observed here became apparent over left frontal electrodes around 1000ms after stimulus presentation, which is at a relatively late stage in the trial.

**Figure 5.**
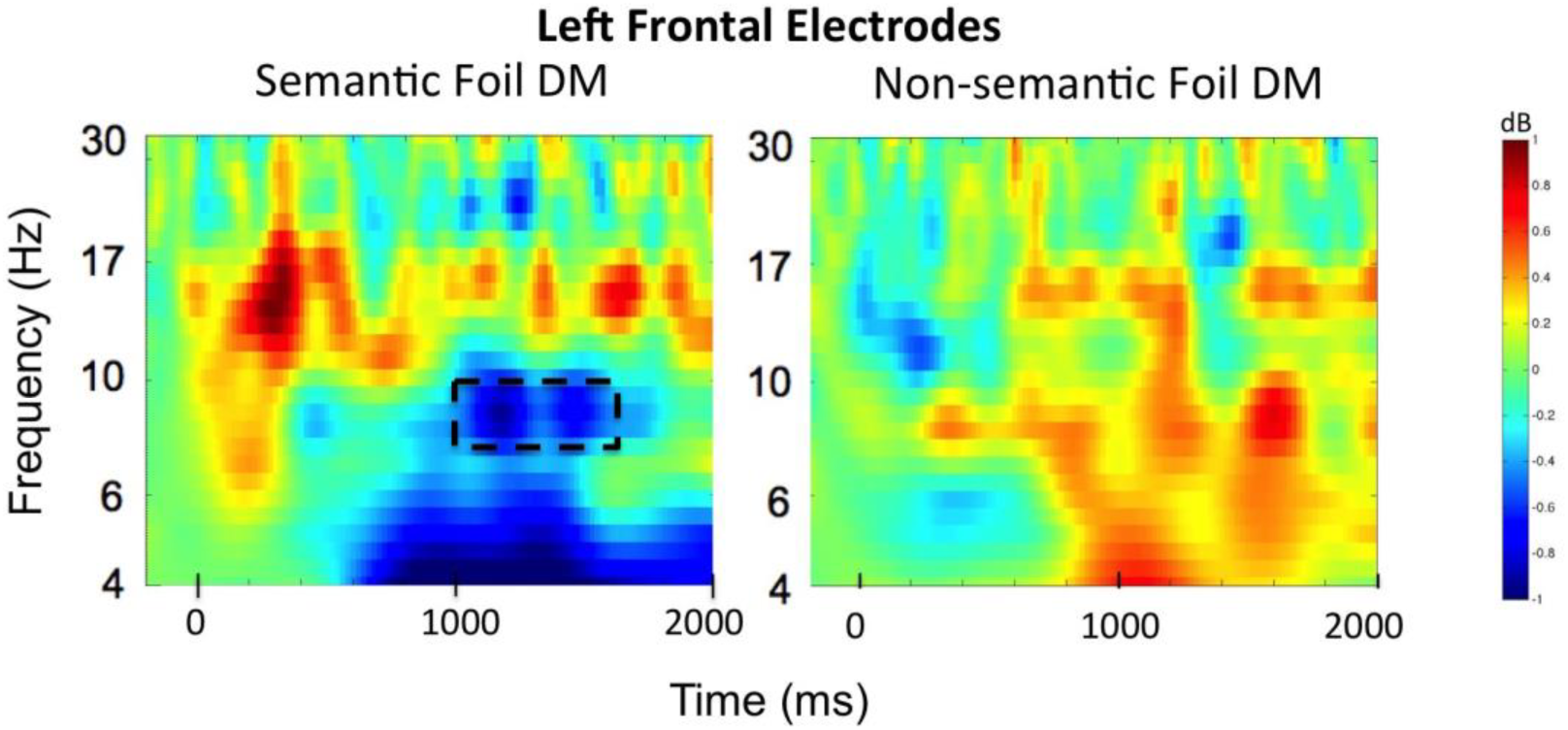
Time-frequency plots from the left frontal cluster illustrating semantic and non-semantic foil subsequent memory (DM) EEG oscillation effects (remembered – forgotten). Successful encoding of semantic foils was uniquely associated with a left frontal alpha power decrease.

**Figure 6.**
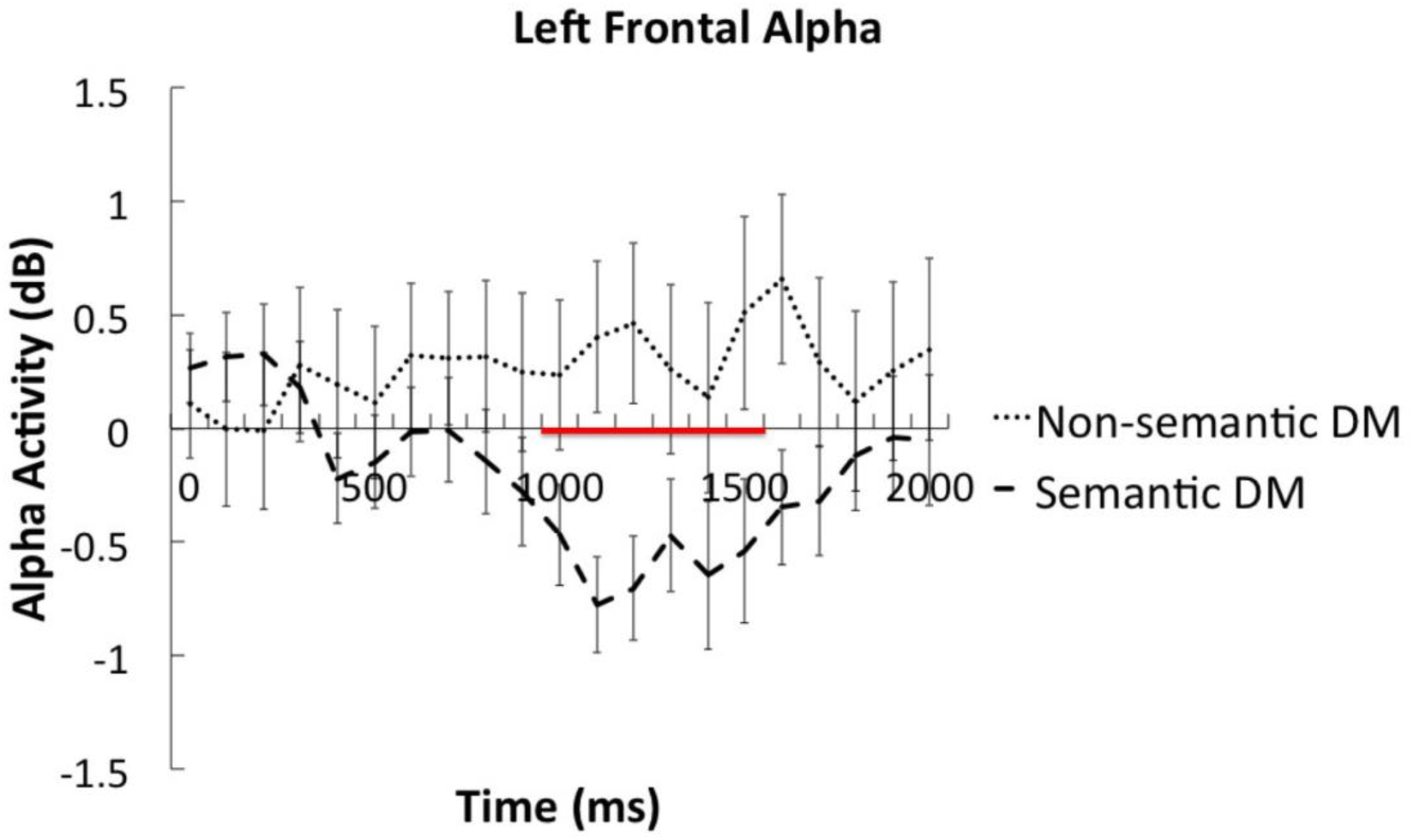
Time courses of left frontal alpha (average of 8–10Hz) differences associated with successful encoding of semantic versus non-semantic foils. Red line on x-axis represents significant time window.

### Relationship Between Phase 1 and Phase 2 Alpha Frequencies

The third part of the analysis assessed the relationship between alpha decreases in phase 1 during semantic processing and alpha decreases in phase 2 during semantic foil recognition, to test whether these effects were functionally related, which would support the hypothesis that the neurocognitive processes engaged during initial encoding are re-implemented when attempting to retrieve information. To examine the relationship between phase 1 and phase 2 alpha activity, we extracted the mean alpha signal that showed a significant effect in the phase 1 (600–1000ms) and phase 2 (1000–1600ms) time windows for each subject and conducted an across-subject correlation. This analysis consisted of two correlations: 1) phase 1 semantic processing alpha activity between 600–1000ms with phase 2 alpha activity in the 1000–1600ms time window associated with semantic foils later remembered; 2) phase 1 semantic processing alpha activity between 600–1000ms with phase 2 alpha activity in the 1000–1600ms time window associated with semantic foils later forgotten.

There was a significant correlation (r = 0.36, p = 0.037; two-tailed) between phase 1 alpha activity associated with semantic processing and phase 2 alpha activity associated with semantic foils later remembered. No such significant correlation (r = 0.17, p = 0.34) was observed between phase 1 semantic alpha related activity and phase 2 alpha activity associated with semantic foils later forgotten (see Figure 7). However, visual inspection of the plots identified an outlier who was more than three standard deviations from the phase 1 semantic alpha activity mean. After removing this outlier, the correlation between phase 1 semantic alpha and phase 2 semantic foils remembered alpha activity became larger (r = 0.54, p = 0.001), but this removal had little effect on the semantic foil forgotten alpha activity correlation (r = 0.23, p = 0.20). The difference between these correlations was marginally significant (t(31) = 1.81, p = 0.08, two-tailed). The results of the correlation analysis are presented in Figure 7.

**Figure 7.**
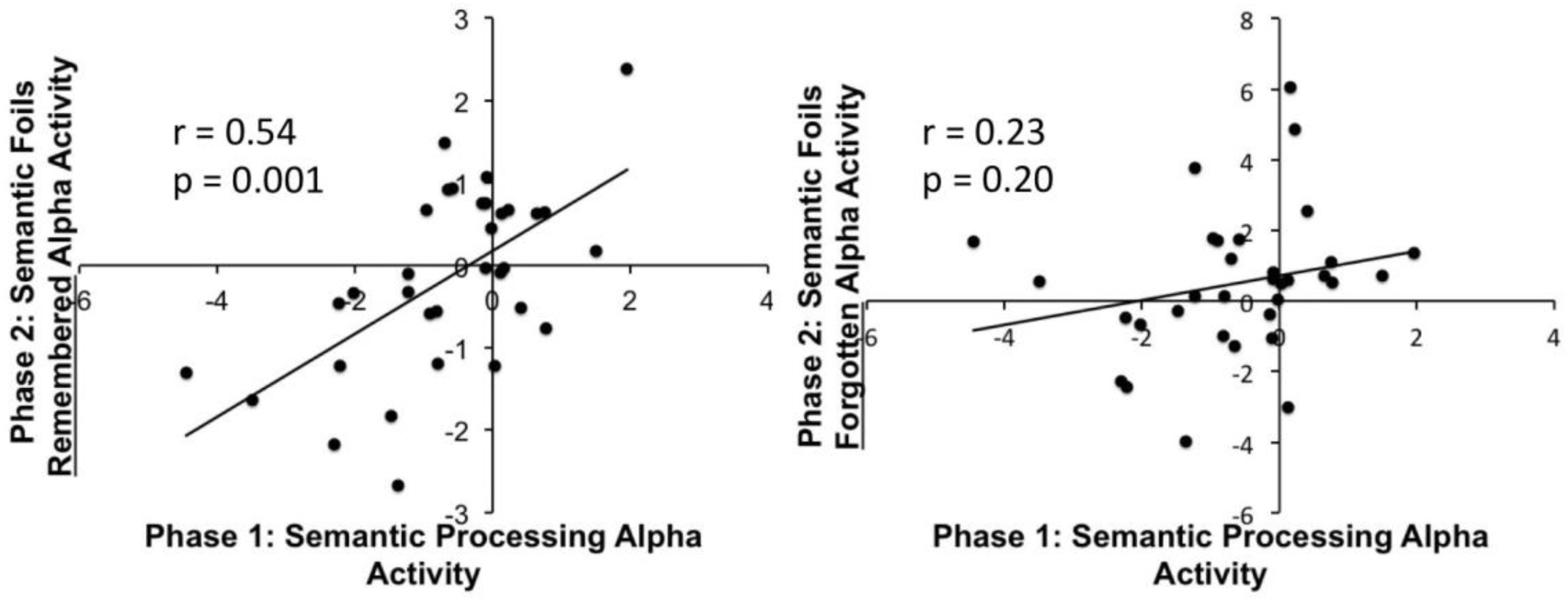
Correlations between phase 1 (600–1000ms) and phase 2 (1000–1600ms) alpha activity.

### Relationship between Alpha Frequencies and Subsequent Foil Recognition

Finally, we examined whether individual differences in phase 2 alpha activity associated with semantic foil encoding correlated with individual differences in behavioral semantic foil recognition in phase 3, which would provide additional evidence that alpha power is functionally related to semantic encoding success. We used the phase 2 alpha power from the 1000–1600ms time window associated with semantic foils that were later remembered and forgotten and correlated this with phase 3 semantic foil recognition accuracy (proportion of correct responses). A significant negative correlation was observed between phase 2 alpha power associated with later remembered semantic foils and phase 3 semantic foil recognition accuracy (r = −0.52, p = 0.002), indicating that individuals who showed the largest alpha power decreases for later remembered semantic foils during phase 2 also were more likely to later recognize semantic foils on the final test. No such relationship was observed for the phase 2 alpha power associated with later forgotten semantic foils and phase 3 semantic foil recognition accuracy (r = −0.04, p = 0.83). The difference between these two correlations was significant: t(31) = −2.82, p = 0.008, two-tailed). Note that removing the outlier that was detected in the previous analysis did not change these results.

**Figure 8.**
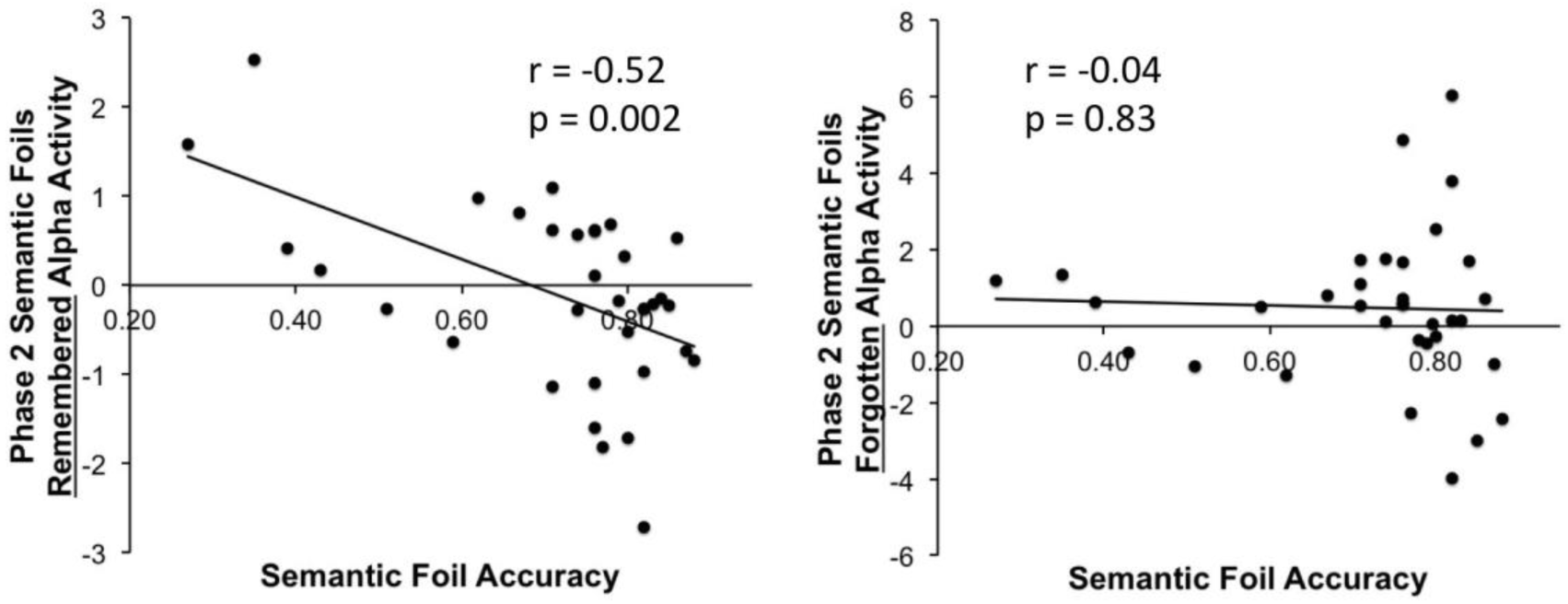
Correlations between phase 2 alpha related activity for both remembered (left) and forgotten (right) semantic foils (1000–1600ms) and subsequent semantic foil recognition accuracy (proportion correct responses).

## Discussion

The aim of the current experiment was to investigate the neural oscillations involved in the successful encoding of new “foil” information presented during a recognition test when participants engage a semantic versus non-semantic processing mode. We tested the hypothesis that attempting to retrieve information from memory involves the re-implementation of the neurocognitive processes that were engaged during initial encoding (Bergström et al., 2015; Jacoby et al., 2005a; Jacoby et al., 2005b; Vogelsang et al., 2016). More specifically, we used the excellent temporal resolution of EEG to examine the temporal dynamics of the encoding of foils to obtain a better understanding of when re-implementation processes occur. It has previously been suggested that the foil effect can be explained by source constrained retrieval processes that re-implement encoding processes in the early stage of a memory test trial to guide memory search as a form of “front-end control” (Jacoby et al, 2005a; Kanter & Lindsday, 2013), predicting that the neural activity associated with such reimplementation should be apparent shortly after a recognition cue is presented. An alternative, though not mutually exclusive, proposal is that control processes may be engaged at a later stage of processing, for example to monitor whether retrieved information is correct (Jacoby et al., 1999; Halamish et al., 2012), or to elicit recollection if initial unconstrained retrieval attempts are unsuccessful as a “late correction” or “back end control” strategy (Jacoby et al., 1999).

Our behavioral findings replicated earlier studies in demonstrating the typical foil effect: Semantic foils were remembered significantly more accurately than non-semantic foils, supporting the idea that participants implemented a semantic processing mode during the semantic memory test (Alban & Kelley, 2012; Danckert et al., 2011; Halamish et al., 2012; Jacoby et al., 2005a; 2005b; Kantner & Lindsay, 2013; Marsh et al., 2009; Vogelsang et al., 2016). Time-frequency analysis of EEG data collected during the initial study phase revealed a power decrease in alpha frequencies over left frontal electrodes between 600–1000ms (and mid/right posterior electrodes between 600–1600ms) during the semantic as opposed to non-semantic task, consistent with prior literature highlighting a role for alpha oscillations in semantic processing (e.g. Bastiaansen et al., 2005; for review see Klimesch, 1999). Importantly, the EEG data from the first recognition test revealed decreases in alpha oscillatory activity in the left frontal electrode cluster between 1000–1600ms that predicted subsequent recognition of semantic, but not non-semantic, foils during the final surprise recognition test. Thus, similar oscillatory activity was associated with semantic processing during initial study and during foil encoding whilst participants were trying to retrieve semantic information. Furthermore, individual differences in alpha activity during the semantic study phase (phase 1) were significantly correlated with individual differences in alpha activity for successfully encoded foils during the semantic recognition test (phase 2), suggesting that the semantic neurocognitive processes that were engaged during initial study were re-implemented during the encoding of foils during the phase 2 recognition test. Finally, we observed that alpha decreases in phase semantic foil encoding during phase 2 were negatively correlated with behavioral semantic foil recognition during phase 3. This result indicates that the larger the decrease in alpha power the better the subsequent recognition memory for semantic foils, suggesting that alpha power is functionally related to semantic encoding success.

Interestingly, alpha power decreases associated with semantic foil encoding became apparent after the average time when participants provided their response, at about 900ms after stimulus presentation, suggesting that alpha oscillations may reflect an implementation of encoding operations at a relatively late processing stage, contrary to what would be predicted if such reinstatement was conducted as part of a front-end control strategy (Gray & Gallo, 2015; Jacoby et al., 2005a; 2005b). In a recent fMRI study, we found that the left inferior frontal gyrus (LIFG) was significantly more active during successful encoding of semantic as opposed to non-semantic foils (Vogelsang et al., 2016). The LIFG has been widely associated with semantic processing (Poldrack et al., 1999; Wagner et al., 1998), but the low temporal resolution of fMRI precluded us from determining whether LIFG activation reflected mentally re-enacting a semantic processing mode early or later in the trial. The timing of the current EEG results suggest that the left frontal alpha decreases, which we tentatively interpret as possibly generated by the LIFG (Vogelsang et al., 2016), may reflect strategic processes that are engaged during a later decision stage of retrieval which facilitates the incidental semantic encoding of foils.

Why did the neural markers of semantic encoding of foils occur so late? Since reinstating encoding operations is an effortful, self-initiated process (Alban & Kelley, 2012), it is possible that participants chose to engage such a strategy in order to elicit recollection only if an initial unconstrained retrieval attempt was unsuccessful. A related account suggests that participants may reinstate encoding operations to verify and possibly correct their initial more automatic retrieval assessments, and such a monitoring strategy may contribute to enhanced encoding of semantic foils together with earlier “front-end” control processes (Halamish et al., 2012). According to Jacoby et al. (1999), participants engage in such late correction strategies primarily when the retrieved information is vague or ambiguous (perhaps eliciting a sense of familiarity without recollection of decisive contextual information). The current oscillatory findings are consistent with reinstatement of encoding operations occurring at a late retrieval stage, but do not rule out the possibility that encoding operations were also reinstated to constrain retrieval at the front-end without being reflected in our EEG results (EEG oscillations of course only capture certain aspects of neural activity).

Our oscillatory findings are consistent with prior literature highlighting a role for alpha frequencies in successful semantic encoding (Hanslmayr et al., 2009; Hanslmayr & Staudigl, 2014; Zion-Golumbic et al., 2009) and semantic processing (Klimesch et al., 2006; Long et al., 2014). In the oscillations literature, alpha frequencies have been linked with a wide variety of cognitive functions ranging from inhibitory processes during memory suppression (Park et al., 2014), to fine-grained resolution of visual processing (Samaha & Postle, 2015), working memory (Sauseng et al., 2009, Myers et al., 2014), and active inhibition of a not-to-be applied rule (Buschman et al., 2012). One of the first studies that found a relationship between decreases in alpha and later memory performance was conducted by Klimesch (1997), who observed that decreases in alpha frequencies over parietal electrodes during semantic encoding were positively correlated with later memory retrieval. Hanslmayr et al. (2009) contrasted deep semantic encoding with shallow non-semantic encoding, and found power decreases in alpha (and beta) frequency bands that were related to successful semantic encoding only and Fellner et al. (2013) showed that alpha likely reflects semantic processing specifically, rather than elaborative and efficient encoding strategies in general. In our experiment, individual differences in alpha power decreases during the semantic recognition test for foils that were later remembered correlated significantly with individual differences in semantic foil recognition accuracy during the final surprise memory test. Together with the subsequent memory alpha effects, these individual differences provide additional converging evidence that alpha power decreases reflect successful semantic encoding.

Jacoby and colleagues (2005a, 2005b) have hypothesized that a possible explanation for the enhanced encoding of semantic versus non-semantic foils in the memory for foils paradigm might lie in the Transfer Appropriate Processing framework and the related Encoding Specificity Principle, both of which predict that retrieval success depends on the amount of overlap between encoding and retrieval processes (Morris et al., 1977; Roediger, 1990; Tulving & Thompson, 1973). While attempting to retrieve words that had either been semantically or non-semantically encoded, participants may mentally re-enact the original study task, resulting in all recognition probes (both old items and foils) being processed semantically during the semantic test block and non-semantically during the non-semantic test block. Semantic retrieval attempts might involve thinking about the meaning of a foil word (e.g. “do I think a strawberry is pleasant?”), whereas non-semantic retrieval attempts might involve examining the letters of the word in the hope that such a strategy will help to decide whether the word is old or new. Such re-enactment may be a relatively late strategy that participants engage in after an initial heuristic familiarity assessment, and may therefore be expressed in neural activity around the time or even after participants have made their memory judgment.

The current time-frequency results in combination with previous research indicate that neural oscillations are a useful tool for studying the temporal dynamics of encoding retrieval overlap (Jafarpour et al., 2014; Staresina et al., 2016; Staudigl & Hanslmayr, 2013; Staudigl et al., 2015; Waldhauser et al., 2012; Waldhauser et al., 2016). Burke et al. (2013), for example, found that high gamma activity (44–100Hz) during successful encoding of information is also observed in similar brain areas during the memory test phase in which previously studied items need to be recalled. Waldhauser and colleagues (2012) observed that decreases in alpha/beta frequencies during retrieval were associated with reactivation of encoded target information, whereas increases in alpha/beta power were associated with the inhibition of encoded distracter information. Cortical reinstatement has also been identified in an entrainment study in which participants studied words presented on flickering backgrounds of either 6 or 10 Hz (Wimber et al., 2012). EEG measurements during successful retrieval of studied words exhibited 6 and 10 Hz frequency oscillations similar to the background flicker rates in which the words had been studied and the strength of this reactivation was related to whether a word was remembered or forgotten (Wimber et al., 2012). More work needs to be done, however, to examine what mechanisms underlie the principle of encoding reimplementation and how that facilitates retrieval. One prominent view is that during retrieval, a cue reactivates only a part of the encoded memory, and that activity of a fraction of the original pattern triggers the reactivation of the entire trace (Rugg et al., 2008). This “pattern completion” process has been linked with the hippocampus and a role for gamma power increases and alpha power decreases has been proposed as a neural mechanism underlying pattern completion (Staresina et al., 2016). However, there is also evidence which suggests that alpha and beta frequency bands in the cortex are important for content specific processing (Hanslmayr et al. 2016), which is in line with our current findings of alpha frequencies representing semantic processing.

To conclude, we investigated the neural oscillations involved in the encoding of new “foil” information presented during a retrieval test as a function of whether the test required participants to retrieve semantic versus non-semantic information. Our findings show that semantic encoding during retrieval attempts was associated with power decreases in left frontal alpha oscillations, which may originate from the LIFG (Vogelsang, et al., 2016). Consistent with previous findings, our results support the view that participants re-implement the distinct neurocognitive operations that were engaged during initial encoding, and we extend previous research by identifying that the time-course of this reimplementation may be at a relatively late processing stage. In this way, memory retrieval can be considered an encoding event, determining whether information will be remembered in the future.

## Acknowledgements

This study was supported by a James S. McDonnell Foundation Scholar Award to JSS, and was carried out within the University of Cambridge Behavioural and Clinical Neuroscience Institute, funded by a joint award from the Medical Research Council and the Wellcome Trust.

## Conflict of interest

None

